# Auditory and semantic cues facilitate decoding of visual object category in MEG

**DOI:** 10.1101/598516

**Authors:** Talia Brandman, Chiara Avancini, Olga Leticevscaia, Marius V. Peelen

## Abstract

Sounds (e.g., barking) help us to visually identify objects (e.g., a dog) that are distant or ambiguous. While neuroimaging studies have revealed neuroanatomical sites of audiovisual interactions, little is known about the time-course by which sounds facilitate visual object processing. Here we used magnetoencephalography (MEG) to reveal the time-course of the facilitatory influence of natural sounds (e.g., barking) on visual object processing, and compared this to the facilitatory influence of spoken words (e.g., “dog”). Participants viewed images of blurred objects preceded by a task-irrelevant natural sound, a spoken word, or uninformative noise. A classifier was trained to discriminate multivariate sensor patterns evoked by animate and inanimate intact objects with no sounds, presented in a separate experiment, and tested on sensor patterns evoked by the blurred objects in the three auditory conditions. Results revealed that both sounds and words, relative to uninformative noise, significantly facilitated visual object category decoding between 300-500 ms after visual onset. We found no evidence for earlier facilitation by sounds than by words. These findings provide evidence for a semantic route of facilitation by both natural sounds and spoken words, whereby the auditory input first activates semantic object representations, which then modulate the visual processing of objects.

## Introduction

In daily life, we continuously integrate information about objects in our environment across sensory modalities. For example, when a distant or occluded object is difficult to identify by the way it looks, we may recognize it by what it sounds like. Indeed, previous studies have demonstrated that audiovisual integration can reshape our perception, causing us to “hear” what we see and to “see” what we hear (McGurk and MacDonald 1976; Spence 2011). When integrating audiovisual information, we rely on the congruence between auditory and visual cues in terms of their object identity, their spatial location, or their temporal synchrony (Noppeney and Lee 2018). To date, while previous studies have used functional magnetic resonance imaging (fMRI) to investigate the anatomical sites of audiovisual interactions (Gau and Noppeney 2016; Werner and Noppeney 2010), it is still unclear how congruent auditory input modulates the time course of visual object processing. Here we used multivariate analysis of magnetoencephalography (MEG) data to probe the temporal dynamics of auditory influences on visual processing of objects that matched in identity. Particularly, we asked how natural sounds and spoken words shape the emergences of visual representations of the same object category, along the time-course of the neural response.

Behavioral studies have shown that auditory cues influence our perception of visual objects in a variety of tasks, including object detection, identification, categorization and visual search (Chen and Spence 2011; 2018; Edmiston and Lupyan 2015; Iordanescu and others 2010; Kim and others 2014; Vallet and others 2013; Weatherford and others 2015). To achieve these effects on visual perception, auditory information could theoretically engage one of two possible routes of facilitation (Figure 1A). One is a semantic route, by which the sound (e.g., the sound of a dog barking) is first interpreted and transformed into a semantic representation (“dog”) before interacting with visual object processing. The alternative is a direct route, by which sounds automatically activate sound-associated object representations in visual cortex, bypassing semantic tagging. Direct auditory-visual mappings could arise out of the natural co-occurrence of visual objects with their corresponding sounds, such as when we see a dog barking. This coupling is consistent; when we hear a bark we see (or expect to see) a dog. Words, on the other hand, do not necessarily co-occur with their corresponding visual object: hearing the word dog does not necessarily mean that we are seeing one.

**Figure 1:**
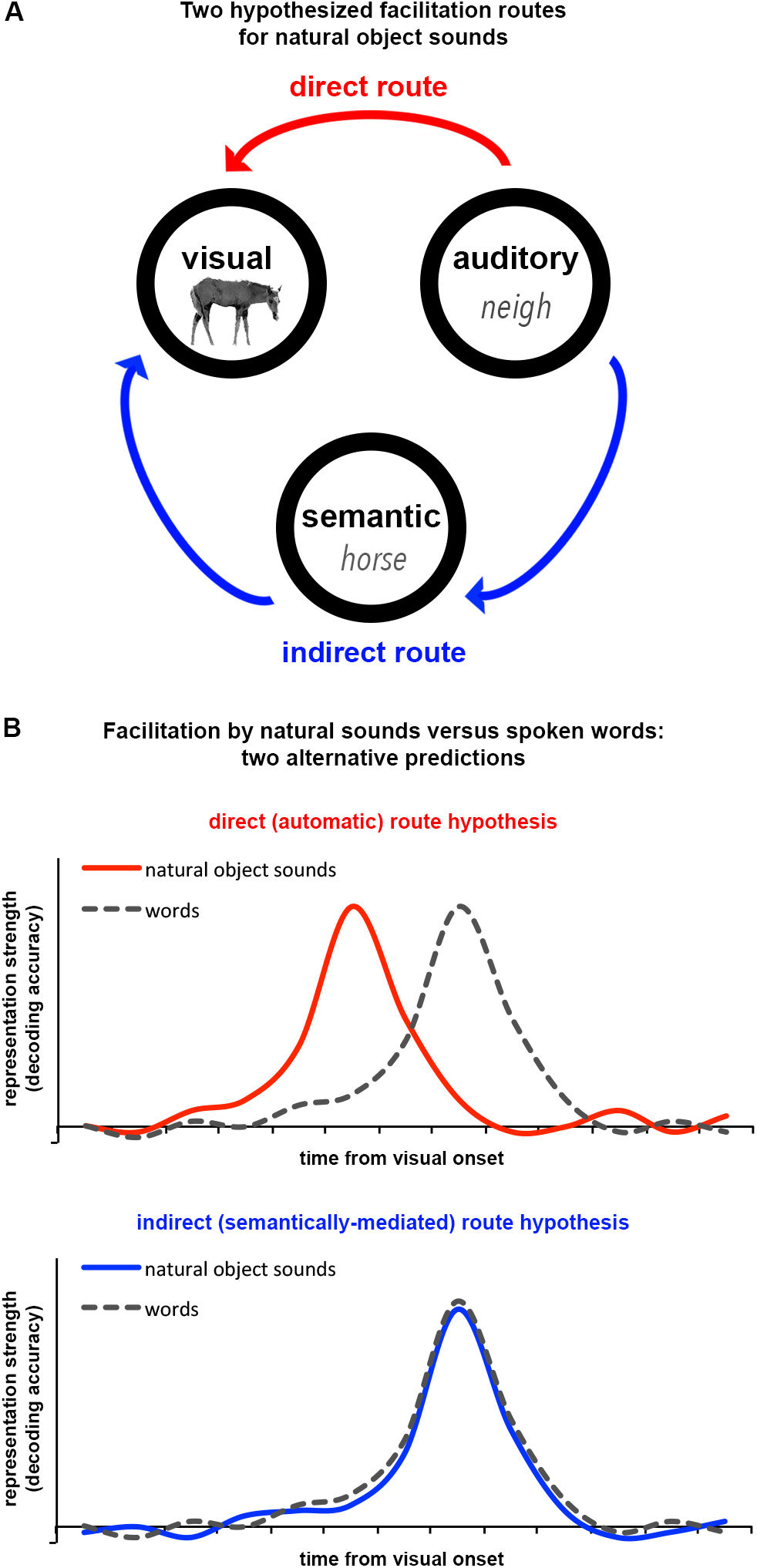
*Two models of audiovisual facilitation*. (A) Two routes of facilitation are hypothesized for naturalistic object sounds; via direct and automatic access to visual representations (in red), or via semantic identification (in blue); (B) Two alternative time-courses are predicted for object representations facilitated by naturalistic object sounds as compared to words, corresponding to the two hypothesized routes in Figure 1A: faster facilitation by sounds than words, corresponding to the direct route (in red), or equally fast facilitation by sounds and words, corresponding to the indirect route (in blue).

In the present study we aimed to test these accounts by comparing the time course of visual object processing facilitated by naturalistic sounds with the time course of visual object processing facilitated by semantic cues (spoken words), both relative to uninformative noise. We hypothesized that if sounds facilitate visual object processing directly and automatically, this modulation would be observed earlier than modulation by word cues. Indirect support for this prediction is found in a series of behavioral studies by Chen and Spence (2011), in which participants were more likely to detect and identify a briefly presented visual object when it was preceded by a congruent sound rather than an incongruent sound. Importantly, detection was more strongly facilitated by naturalistic sounds than by spoken words (see also Chen and Spence 2018). Since pictures are matched more rapidly with naturalistic sounds than with words (Saygin and others 2005), it was proposed that naturalistic sounds may directly affect object representations, whereas words may do so via a longer route (Chen and Spence 2011).

Further evidence for (anatomically) early audiovisual interaction comes from recent neuroimaging studies showing that auditory cues influence activity in early visual areas (for review see Murray and others 2016) and vice versa (Werner and Noppeney 2010). For example, Vetter and others (2014) demonstrated that it is possible to decode the category of natural sounds based on patterns of the fMRI signal in V1. In another fMRI study, de Haas and others (2013) found that incongruent sounds significantly impaired visual stimulus decoding from area V2. Investigating the time course of facilitation, electrophysiological studies using artificially-learned audiovisual associations (e.g. an oval and a tone; Cappe and others 2010; Fort and others 2002; Giard and Peronnet 1999; Molholm and others 2002), reported audiovisual interaction within the first 200 ms of response. Finally, transcranial magnetic stimulation (TMS) findings suggest that sounds may increase early visual cortex excitability (Romei and others 2009). Altogether, these studies provide some evidence that auditory cues may affect early stages of visual processing, and not only higher-order regions of audiovisual integration (e.g. the superior temporal sulcus; Beauchamp 2005; Watson and others 2014). Yet the time-course for audiovisual facilitation of naturalistic objects remains largely unknown, and no study has directly compared facilitation by sounds with facilitation by words.

In the current study, combining a new approach to multisensory facilitation with MEG multivariate analysis allowed us to directly probe the temporal dynamics of audiovisual facilitation encoded in the neural signal. Thereby, we examined the effects of auditory cues on the neural representations of animate and inanimate visual objects. Previous electrophysiological studies have shown that visual object animacy is represented in the multivariate response pattern across the scalp, emerging as early as 80 ms after visual onset (Carlson and others 2013; Cichy and others 2014). Furthermore, we have recently shown that after 300 ms, degraded object-animacy representations rely not only on feed-forward visual object information, but also on external cues such as the background scene (Brandman and Peelen 2017). We used a similar approach to first measure the time-course of multivariate representation of visual object animacy in degraded objects, along the MEG signal. Second, extending this approach to audiovisual integration, we examined the facilitatory effects of natural sounds and spoken words on the time-course of visual representation. We hypothesized that if naturalistic object sounds automatically activate object representations in the visual system, bypassing semantic interpretation, then they would facilitate visual processing of objects earlier than words. Alternatively, if natural sounds take an indirect semantically-meditated route, facilitation would be as late as with words (Figure 1B).

## Materials & Methods

We measured the multivariate representations of object animacy in MEG signals evoked by degraded objects with sounds, words and noise, and by gray rectangles with sounds, while participants performed a 1-back repetition detection task. In a separate intact-objects experiment used for classifier training, participants viewed intact animate and inanimate objects, fully visible and in high resolution, presented with no auditory input. We used cross-decoding classification of object category (animate/inanimate) to compare the multivariate response patterns evoked by intact objects to those evoked by the audiovisual experiment stimuli. This allowed us to isolate the contribution of each type of auditory cue (sound, word, noise) to the visually-evoked neural representation of object animacy, in order to characterize their temporal dynamics of facilitation.

All procedures were approved by the ethics committee of the University of Trento.

Stimuli and analysis code are available at: https://github.com/tbrandman/stimuli_and_analysis_code_BrandmanEtAl2019

### Participants

Twenty-five healthy participants (8 male, mean 25 years ± 4.8 SD) were included. All participants had normal hearing, and normal or corrected to normal vision, and gave informed consent. Sample size was chosen to match that of our previous study using similar MEG decoding methods (Brandman and Peelen 2017). One additional participant was excluded due to misunderstanding the task. Three additional participants were excluded from data analysis due to excessive signal noise.

### Stimuli

Eighty-four photographs of objects, 42 animate and 42 inanimate, were cropped, transformed to grayscale, blurred and presented on light gray background (see Figure 2A for examples). For the visual uninformative-control condition, a uniform gray rectangle (of the same dimensions) was created using the mean luminance corresponding to each of the 84 object stimuli. Eighty-four natural sounds, matched to each of the 84 visual objects (e.g. barking dog, alarm clock ringing, etc.), were cropped to 1500 ms length and were the same in right and left channels. For the auditory uninformative-control condition, noise tracks were generated from each of the 84 natural sounds, by scrambling all sampling points. Word stimuli (in Italian) describing each of the 84 visual objects were recorded in the lab by a native Italian speaker. Similar to the natural sounds, words were under 1500 ms long and were the same in the right and left channels. To avoid familiarity effects passing between the three auditory conditions (natural sounds, words, noise), the stimulus set was split into three, such that different pictures were presented for objects with sounds, objects with words and objects with noise within a given subject. The rectangles with sounds matched the objects with noise. The three sets were counterbalanced across subjects. For example, for stimuli bird1, bird2 and bird3, one subject would view bird1 with sound, bird2 with word and bird3 with noise, and also the gray rectangle corresponding to bird3 with sound, while a second subject would view bird1 with word, bird 2 with noise and corresponding gray rectangle with sound, and bird 3 with sound, and so on. The intact-objects experiment included the 84 objects from the audiovisual experiment (“old”) and an additional set of 84 new objects (“new”) that were matched for category and subcategory of the audiovisual experiment set. The new exemplars were added in order to prevent over-fitting of the classifier to lower-level visual features of the audiovisual object exemplars. Image processing was the same as in the audiovisual experiment, excluding the blurring step, such that objects were presented intact. Object photographs were obtained from previous stimulus sets (Downing and others 2006; Konkle and Oliva 2012) and open online image databases. All visual stimuli and fixation points were presented centrally at a visual angle of 5.25 degrees (350 × 350 pixels).

**Figure 2:**
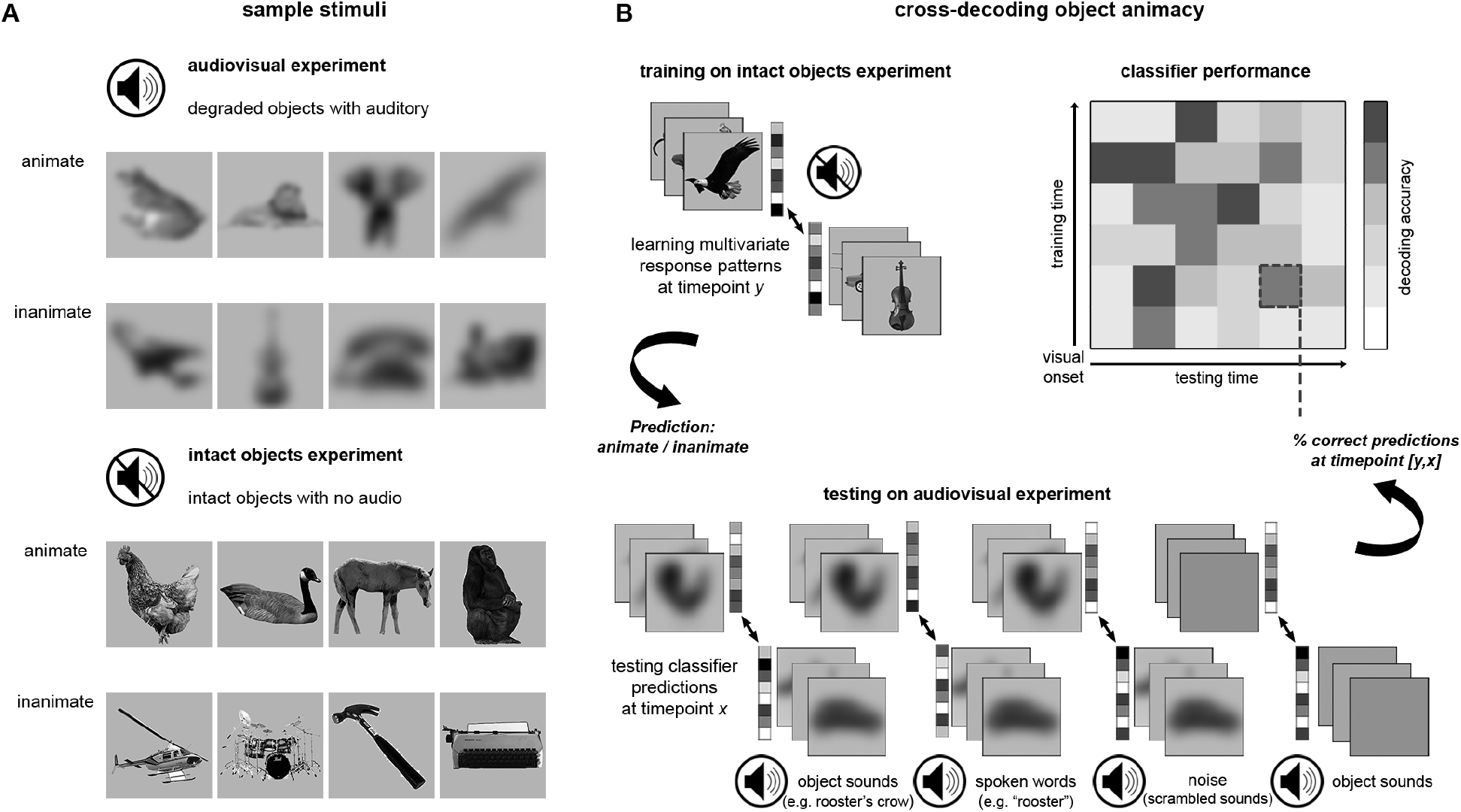
*Experimental design*. (A) Sample visual stimuli. Left: Audiovisual experiment presenting images of animate and inanimate degraded (blurred) objects, preceded by the onset of an auditory cue. Right: Intact-objects experiment presenting images of animate and inanimate intact (high-resolution) objects with no sounds. (Object images taken from open online databases and from Downing and others 2006; Konkle and Oliva 2012); (B) Cross-decoding analysis, whereby a classifier was trained on animate/inanimate intact object discrimination from the response patterns across MEG sensors, and tested on discrimination of animate/inanimate degraded objects with natural sounds, spoken words and noise, and on mean-luminance rectangles with natural sounds. The result is the mean classification accuracy for each of the four main experiment conditions, tested within each time-point separately. At the top right corner we illustrate the time-by-time matrix structure for results presented in Figure 3, in which mean classification accuracies are assigned to the time-point corresponding to their training time (Y axis) and testing time (X axis).

### Procedure

On each trial, participants viewed a single briefly presented (50 ms) visual stimulus. During audiovisual experiment runs, prior to visual onset, participants heard a sound onset between 900 and 1100 ms before visual onset. This was done to allow enough time for the semantic processing of words prior visual onset. Although natural sounds tend to occur more often simultaneously with vision in real life, here we had to equate their priming interval with that of words. In addition, the exact time between auditory onset and visual onset was randomly jittered within the range of 900 to 1100 ms to allow us to separate visual-onset evoked responses from auditory-onset evoked responses. The auditory cue lasted for ~800-1500 ms, in most cases offsetting after visual offset. After sound offset, inter-trial interval (ITI) jittered between 1500-2500 ms. During the intact-objects experiment, no sound was presented, and ITI after visual offset jittered between 1400-2400 ms. In the absence of visual stimuli a central fixation dot was presented on the screen. During all runs, participants performed a 1-back task on the visual stimuli, in which they were asked to press a button whenever the same image was presented twice in a row.

The audiovisual experiment consisted of 6 runs of ~449 s duration, each composed of 5 fixation breaks (8 s each), 5 repeated trials (1-back targets) and 14 trials per condition: animate/inanimate x object-with-sound / object-with-word / object-with-noise / gray-rectangle-with-sound (112 trials/run). The audiovisual experiment was followed by the intact-objects experiment. The intact-objects experiment consisted of 2 runs of ~437 s duration, each composed of 11 fixation breaks (8 s each), 11 repeated trials (1-back targets) and 42 trials per condition: animate/inanimate x old/new (168 trials/run). This resulted in a total of 84 trials per condition across audiovisual experiment runs, and 84 trials per condition across intact-objects experiment runs. Within each run, trials were randomly intermixed.

### Data Acquisition and Preprocessing

Electromagnetic brain activity was recorded using an Elekta Neuromag 306 MEG system, composed of 204 planar gradiometers and 102 magnetometers. Signals were sampled continuously at 1000 Hz and band-pass filtered online between 0.1 and 330Hz. Offline preprocessing was done using the Elekta MaxFilter/MaxMove software, MATLAB (RRID:SCR_001622) and the FieldTrip analysis package (RRID:SCR_004849). Signals were filtered of external noise and spatially realigned with the Maxwell Filter and Move, using signal space separation algorithms (Taulu and Simola, 2006). Data were then demeaned, detrended, down-sampled to 100 Hz and time-locked to visual onset. The data were averaged across trials of the same exemplar across runs, resulting in a total of 112 unique test trials throughout the audiovisual experiment (14 per condition), and 168 unique train/test trials throughout the intact-objects experiment (42 per condition).

### Multivariate Analysis

Multivariate analysis was performed using CoSMoMVPA toolbox (Oosterhof and others 2016) (RRID:SCR_014519). Analysis followed a similar procedure as in Brandman and Peelen (2017), in which the same cross-decoding approach was used to successfully decode visual object category from degraded object stimuli. Decoding was performed across posterior magnetometers (48 channels) of each participant, between 0 and 600 ms. Prior to decoding, temporal smoothing was applied by averaging across neighboring time-points with a radius of 2 (20 ms). An LDA classifier discriminated between response patterns to animate vs. inanimate objects, using the default regularization value of .01 for covariance matrix. The decoding approach is illustrated in Figure 2B. First, decoding of intact object animacy, in the absence of auditory input, was measured within the intact-objects experiment, by training on old-object trials (i.e. objects included in the audiovisual experiment set), and testing on new-object trials. Next, cross-decoding was achieved by training on the intact-objects experiment, and testing on each of the audiovisual experiment conditions (object-with-sound, object-with-word, object-with-noise, gray-rectangle-with-sound). All intact-object exemplars (old and new) were used in training the cross-decoding classifier, because no differences were found in cross-decoding using only old or only new intact objects in the training dataset. Decoding was performed for every possible combination of training and testing time-points between 0 and 600 ms, resulting in a 60 × 60 matrix of 10 ms time-points, for each of the tested conditions, per subject. In addition, to generate a measure of same-time cross-decoding, decoding accuracy of each time-point along the diagonal of the matrix was averaged with its neighboring time-points at a radius of 2 (20 ms in every direction).

### Significance testing

Significance was tested on contrasts across the time-by-time matrix as well as along same-time cross decoding. Significance was tested for each time-point by computing random-effect temporal-cluster statistics corrected for multiple comparisons. This was accomplished via t-test computation over 1000 permutation iterations, in which the sign of samples was randomly flipped (over all features) after subtracting the mean, and using threshold free cluster enhancement (TFCE) as cluster statistic, with a threshold step of .1. Significance was determined by testing the actual TFCE image against the maximum TFCE scores yielded by the permutation distribution (TFCE, *p* < 0.05) (Smith and Nichols 2009; Stelzer and others 2013). Significant above-chance (50%) decoding of intact objects in the intact-objects experiment was tested across the entire time-by-time matrix. Significant time-points were then used to define a temporal mask for cross-decoding significance testing.

### Behavioral sound recognition test

In order to exclude potentially ambiguous sounds in a post-hoc MEG analysis, sound recognition was measured, in a separate group of subjects, in an online behavioral experiment conducted via Amazon Mechanical Turk. Although the physical experimental settings (e.g. sound quality, room noise) in online data collection vary across participants, these settings did not systematically vary across exemplars because each participant completed trials of all exemplars in random order.

#### Participants

Eighteen participants (11 male, mean 42 years ± 11.7 SD) were included in the behavioral experiment. All participants gave informed consent. Two additional participants were excluded from data analysis due to below-chance performance and/or incompliance with task instructions.

#### Stimuli

Stimuli included all of the natural animate and inanimate sounds used in the MEG experiment. No other auditory or visual stimuli were presented.

#### Procedure

On each trial, participants listened to a sound while viewing a blank screen, and pressed a button at the moment they recognized it. They then typed in what they thought was the object that had made the sound. Finally, they rated how clearly the sound had conveyed the object, on a scale from 1-very ambiguous to 8-very clear. Each sound was presented until response, once per subject, in random order, resulting in 84 test trials. Four practice trials preceded the test trials, using sounds that had not been included in the MEG experiment.

#### Response Quantification

Sound recognition was quantified by coding open answers into 50 fine-grain object categories, including each type of animal or inanimate object. A single response was defined as correct if the answer fell within the fine-grain category assigned to the exemplar, and incorrect for any other response. For each sound, group recognition accuracy was calculated as the ratio of correct responses across subjects. Recognition response times and clarity ratings were highly correlated with recognition accuracy (r > 0.5) and were not further analyzed.

#### Post-hoc MEG trial exclusion

In a post-hoc analysis of MEG responses evoked by objects with natural sounds, we excluded sounds that were accurately recognized by less than 50% of participants in the behavioral experiment. This resulted in 11 excluded inanimate sounds and 2 excluded animate sounds. To balance the number of sounds between animate and inanimate condition, we further excluded the next 9 worst-recognized animate sounds. Thus, the total number of excluded sounds was 22 (11 animate and 11 inanimate). Cross-decoding using the remaining 62 well-recognized sounds was performed in the same way as in the main analysis, and is reported under “objects with clear sounds”, separately from objects with (all) sounds, in the final section of the results.

## Results

### Decoding the animacy of intact visual objects

In a first analysis, we assessed the temporal dynamics of animacy decoding of intact objects within the intact-objects experiment. Classifiers were trained and tested on 10 ms time intervals from 0 ms to 600 ms relative to visual onset, resulting in a 60×60 time-by-time matrix of decoding accuracy. Results showed significant decoding of object animacy along the diagonal of the matrix (same training and testing time-points), peaking around 200 ms after visual onset (Figure 3A). These results replicate previous findings of animacy decoding using MEG (Brandman and Peelen 2017; Carlson and others 2013; Cichy and others 2014).

**Figure 3:**
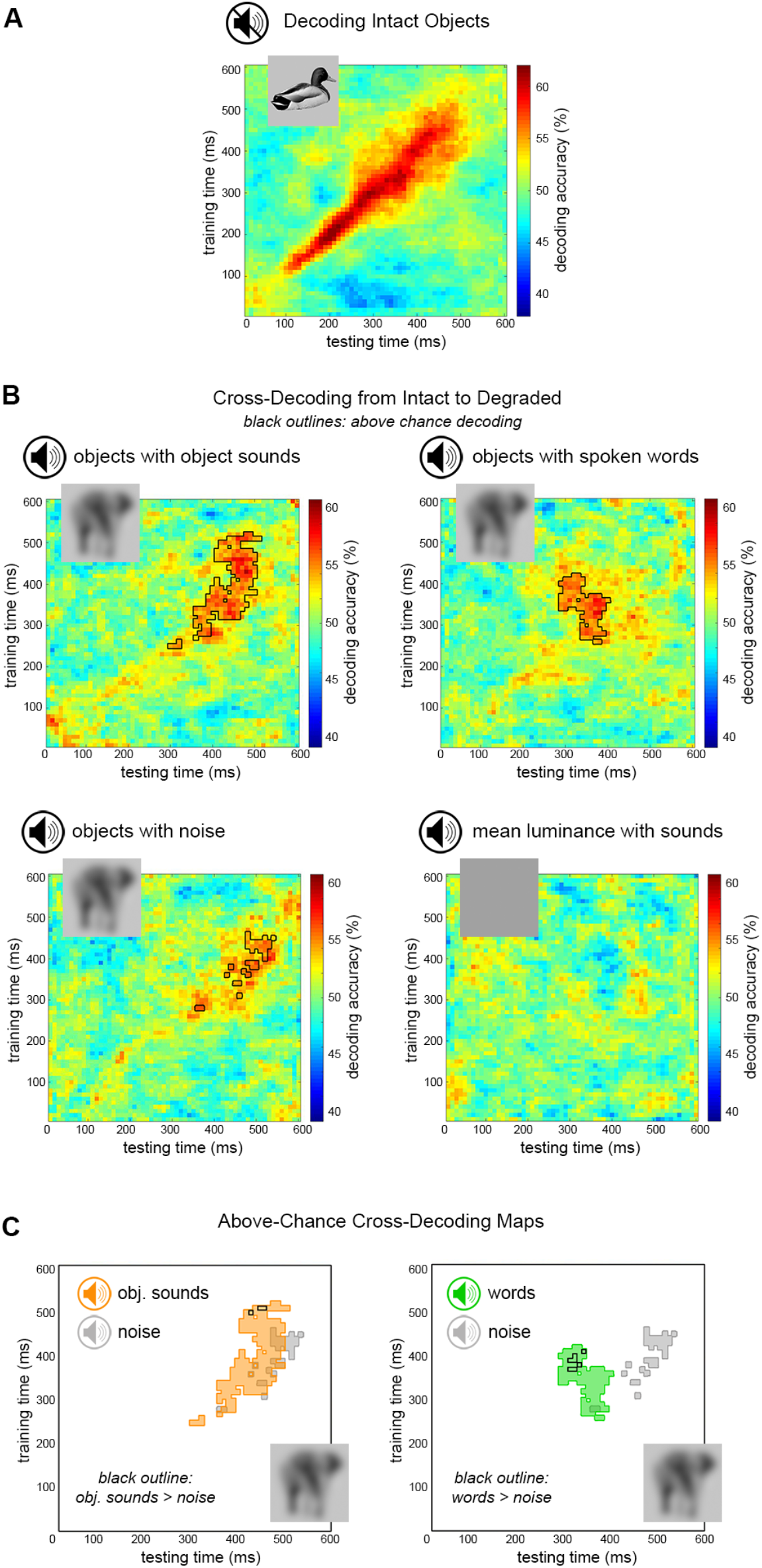
*MEG decoding of object animacy*. Matrices present decoding results in a time-by-time space from visual onset to 600 ms. (A) Decoding accuracy of intact object animacy, averaged across subjects. Object animacy was successfully decoded from intact objects with no sounds, peaking around 200 ms. (B) Cross-decoding accuracy of degraded object animacy, averaged across subjects. Object animacy was decoded above chance (black outline; *TFCE 1-tail p* < 0.05) from degraded objects with sounds (top left), words (top right) and noise (bottom left), but not from of mean-luminance rectangles with sounds (bottom right). (C) Above-chance cross-decoding clusters. Black outlines represent significantly better decoding of object animacy from degraded objects with informative auditory cues (sounds or words) than with noise (paired; *TFCE 1-tail p* < 0.05). Both sounds (left) and words (right) significantly facilitated the representations of object animacy compared to noise.

### Cross-decoding visual object animacy from audiovisual stimuli

Next, we examined the decoding of object animacy in the audiovisual experiment, using a cross-decoding approach (Figure 2B). Classifiers were trained to discriminate animacy of intact visual objects in the intact-objects experiment, and then tested on animacy discrimination in degraded objects with sounds, degraded objects with words, degraded objects with noise, and gray rectangles with sounds. Decoding accuracies are presented in Figure 3B. Animacy of degraded objects could be reliably decoded (against chance; *TFCE 1-tail p* < 0.05) when presented with each of the auditory stimuli: natural sounds, spoken words, or noise. In contrast, object animacy could not be decoded from gray rectangles with animate and inanimate natural sounds (against chance; *TFCE 1-tail p* > 0.05), showing that auditory information on its own was insufficient in activating visual representations of object animacy.

### Facilitating effects of informative versus uninformative sounds

Above-chance decoding across the time-by-time matrices is outlined in Figure 3C, showing that degraded objects with sounds, words, and noise are decodable within different, yet partially-overlapping time clusters. To directly test whether sounds and words facilitated decoding relative to noise, we tested their pairwise comparisons within all time-points showing above-chance decoding for any of the audiovisual experiment conditions. Outlined in black over Figure 3C, analyses revealed better decoding of degraded objects with sounds than with noise at multiple time-points between 420-510 ms after visual onset (paired; *TFCE 1-tail p* < 0.05), and better decoding of degraded objects with words than with noise at multiple time-points between 300-410 ms after visual onset (paired; *TFCE 1-tail p* < 0.05). Thus, both natural sounds and spoken words significantly improved decoding of degraded object animacy relative to noise.

Finally, we asked whether words and sounds differ in their speed of facilitation. Above-chance decoding across the time-by-time matrices of the two conditions is overlaid in Figure 4A, showing an earlier above-chance time-cluster for words than for sounds. To directly test the difference in decoding time of degraded objects with sounds versus words, we tested their pairwise comparison within all time-points showing above-chance decoding for any of the audiovisual experiment conditions (including noise). Outlined in black over Figure 4A, analyses revealed better decoding of degraded objects with sounds than with words at multiple time-points between 450-510 ms after visual onset (paired; *TFCE 2-tail p* < 0.05), and better decoding of degraded objects with words than with sounds at multiple time-points between 300-410 ms after visual onset (paired; *TFCE 2-tail p* < 0.05). This shows that sounds and words facilitated degraded objects at different stages along the time-course of visual animacy representation. Particularly, words were faster than sounds in reconstructing representations of animacy evoked by visual objects.

**Figure 4:**
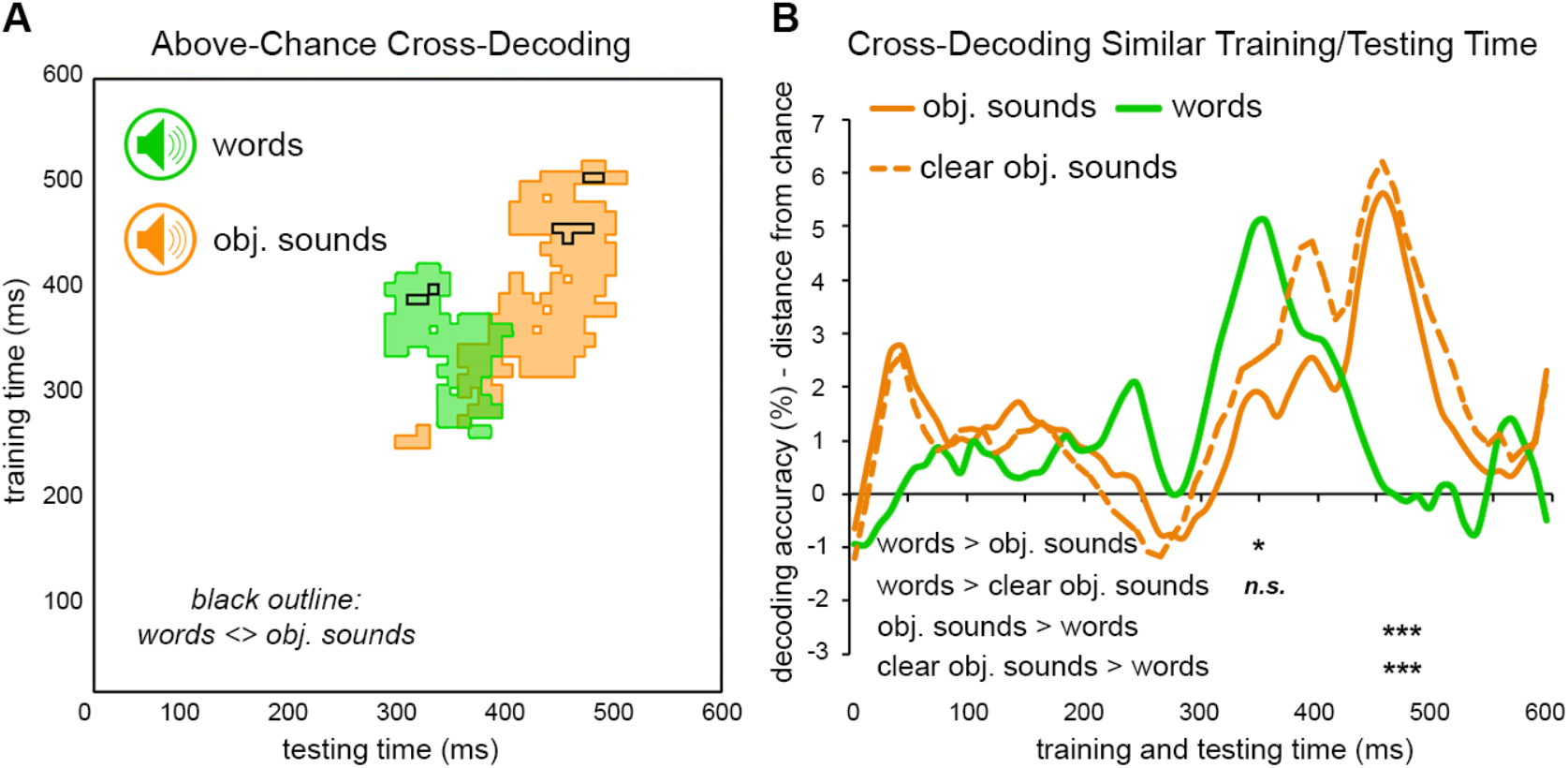
*Facilitation by sounds versus words*. (A) Above-chance cross-decoding clusters. Black outlines represent significantly better decoding of object animacy from degraded objects with sounds than with words and vice versa (paired; *TFCE 2-tail p* < 0.05). Initial analysis including all presented sounds and words revealed an earlier time-window for facilitation by words compared to sounds. (B) Cross-decoding accuracy along the smoothed time diagonal (matched training and testing times). Object animacy was decoded significantly better with words than with sounds, but not better than with clear sounds, at 350 ms. Decoding was better with sounds and clear sounds than with words at 460-480 ms. Data are represented as mean distance from chance (50% decoding accuracy), *paired; *TFCE 2-tail p* < 0.05 for each time-point indicated.

### How does sound interpretability affect the time-course of facilitation?

Referring back to the two models illustrated in Figure 1, our results do not support faster facilitation for sounds relative to words. Rather, facilitation by spoken words was seen even earlier than facilitation by natural sounds, a result that neither of the two models predicted. We therefore asked whether this difference could have been mediated by the efficacy of the facilitator, i.e. by the strength of semantic representation. We hypothesized that if facilitatory effects of natural sounds rely on their translation into semantic representations of animacy, then the ambiguity of some sounds may harm their efficacy and slow down the facilitation. To test this, we conducted an online behavioral experiment, in which a new group of participants were asked to identify the sounds used in the MEG, when presented with no visual input (see Material and Methods). Based on these behavioral data, 22 sounds were marked as relatively ambiguous. We then returned to the MEG data, examining cross-decoding accuracies for degraded objects with sounds, including only trials with clear (unambiguous) sounds.

When testing only clear sounds, the advantage of words over sounds at the earlier time cluster (~300-400 ms; Figure 4A) was no longer significant (paired; *TFCE 2-tail p* > 0.05), whereas the advantage of sounds over words (paired; *TFCE 2-tail p* < 0.05) and over noise (paired; *TFCE 1-tail p* < 0.05) at the later time cluster (~400-500) was preserved. We illustrate the time-courses of facilitation by words, sounds and clear sounds in Figure 4B, by presenting decoding accuracies (minus chance: 50%) along similar training and testing times (±20 ms). Decoding of degraded objects with sounds and with clear sounds peaked at 460 ms after visual onset, where it was also significantly better than decoding of degraded objects with words (paired; *TFCE 2-tail p* < 0.05). Decoding of degraded objects with words peaked at 350 ms after visual onset, where it was significantly better than decoding of degraded objects with sounds (paired; *TFCE 2-tail p* < 0.05), but not clear sounds (paired; *TFCE 2-tail p* > 0.05). In addition, decoding of degraded objects with clear sounds seems to reveal a new peak at 390 ms after visual onset, closer in time to peak word-driven facilitation.

## Discussion

The present study investigated the temporal dynamics by which natural sounds and spoken words facilitate the visual representations of objects, across the MEG multivariate response pattern. Results show that both sounds and words significantly facilitated decoding of degraded object category (animate vs inanimate) relative to uninformative noise, between 300 and 510 ms after visual onset. We found no evidence that natural sounds elicited faster facilitation than words. Particularly, our findings show that both natural sounds and spoken words facilitate the multivariate representation of visual object animacy within the first 500 ms of neural response. In a previous study, we used a similar cross-decoding approach to show facilitation of object animacy decoding by scene context (Brandman and Peelen 2017). In that study, degraded objects were better decoded when embedded within congruent scene context from around 320 ms after stimulus onset. The current results show that the neural representations of visual objects are shaped not only by extrinsic visual information but also by extra-visual, auditory and semantic information. Altogether, our findings characterize the time-course of auditory- and semantic-driven facilitation, showing that natural sounds and spoken words bias the neural representation of visual object category at similar stages of visual processing.

The current results challenge the hypothesis that natural sounds access earlier, or more direct, representations of objects relative to words; our data reveal no temporal advantage of sounds over words in facilitating the neural representations of objects. In fact, initial analysis even showed faster facilitation by spoken words than by natural sounds. However, when restricting the analysis to clear, unambiguous sounds, the temporal advantage of words over sounds was no longer significant. Our findings therefore indicate a similar temporal window for both sounds and words in facilitating visual object animacy. As such, both types of cues may access similar stages of object representation. Particularly, as we find no evidence for faster facilitation for natural sounds over spoken words, we propose that similar to words, natural sounds exert a top-down effect of semantic expectation on the visual representation of objects. Thereby, our data are most in line with an indirect, semantic route for auditory-driven facilitation of visual object representation (Figure 1, in blue).

In the present study, we examined passive (or automatic) facilitation driven by task-irrelevant cues: participants were required to detect image repetition but could ignore the auditory input. Considering the interaction between attention and multisensory integration (Talsma and others 2010), we expect that the time course of auditory-driven facilitation in part depends on the degree to which the auditory cues are task-relevant. For example, when auditory or symbolic cues are used to direct attention to specific object categories, the modulatory effect of these cues on visual processing is observed within the first 200 ms of the MEG response, even for objects embedded in complex natural scenes (Kaiser and others (2016). Task-dependent modulation was also found in an EEG study by Molholm et al., (2004), showing audiovisual congruity effects of objects with naturalistic sounds within the first 200 ms of response, for target objects but not for non-target objects. Similarly, behavioral studies have shown that symbolic category cues can influence early perceptual sensitivity (Lupyan and Ward 2013; Stein and Peelen 2015). In these studies, the simple detection and localization of visual objects was facilitated by written or spoken words. We would thus expect a speeded effect of facilitation, relative to the current findings, by both sounds and words when these cues are used to direct attention to the visual object category. However, in this scenario, sounds and words would engage the same top-down attention mechanism, which would again result in similar facilitation for both types of cues, as in the current study.

Another factor that may affect the speed of facilitation is the specificity of the auditory cue. The sounds used in the present study provided information about the object at the basic level (e.g., horse). It is possible that the use of more specific sound-object pairs would result in earlier facilitation. For example, an image of a horse on its hind legs with its mouth open may be faster facilitated by the sound of a neigh, priming this specific depiction of a horse, than by the sound of a trot, priming a more general concept of a horse. Support for the case of early facilitation in high-specificity paired features can be seen in previous electrophysiological studies (Fort and others 2002; Giard and Peronnet 1999), showing audiovisual integration effects for particular combinations of tones and ovals, within the first 200 ms of response. In addition, fMRI findings (Kok and others 2012) revealed that in early visual cortex, grating orientation was better decoded following a predictive tone, suggesting that expectations facilitate vision by sharpening early visual representations. Furthermore, representations of expected stimuli in the MEG signal emerged even before they were presented (Kok and others 2017). Contextual contingency and expectation were also shown to modulate audiovisual interactions in EEG, by manipulating spatial predictability (Matusz and others 2016). Finally, our own findings show that restricting the analysis to more accurately identified sounds revealed an earlier temporal window for facilitation. Considering these previous studies and our current findings, we propose that visual features may be facilitated by highly specific auditory cues at an early stage of their representation, while representations of visual category are facilitated by category-level cues at a later stage, as in the present study. Future studies could directly test this hypothesis by manipulating the level of audiovisual specificity.

While more specific auditory cues may lead to earlier facilitation, it is important to note that examining a high-tier category representation, i.e. animacy, does not in itself explain the relatively late onset of facilitation reported here. First, extant evidence demonstrates that animacy is not a late distinction in visual object discrimination. For example, in MEG object decoding, animacy was found to emerge at similar times as basic-level categories, and peak only tens of milliseconds later, within the first 250 ms of the neural response (e.g. Carlson and others 2013; Cichy and others 2014). Indeed, when controlling for perceptual differences between animate and inanimate objects, MEG patterns may no longer differentiate between them (Proklova and others 2019). In neuroimaging studies, mid-level visual features were shown to distinguish animacy in visual cortices (Grill-Spector and Weiner 2014), as animate objects differ from inanimate objects in terms of their characteristic shapes and other category-associated visual features (Levin and others 2001; Long and others 2017; Schmidt and others 2017; Zachariou and others 2018). Second, in our intact-object experiment, animacy was decoded above chance as early as ~100 ms, indicating that the animacy classifier was sensitive to visual features distinguishing animacy at pre-semantic stages. Therefore, our finding of late degraded-object representations does not reflect an insensitivity of the animacy classifier at earlier time points. Rather, the delay in the audiovisual condition provides evidence that auditory cues facilitate later representations of visual object animacy.

In sum, we have shown that auditory cues shape the representation of visual object category decoded from the neural response pattern across the scalp. Particularly, both natural sounds and spoken words facilitated object animacy representations between 300 and 500 ms after visual onset. This is consistent with an indirect route for auditory-driven facilitation of visual object representation (Figure 1, in blue), similarly for natural sounds and spoken words. To conclude, our findings provide evidence for a semantic route of visual facilitation by both natural sounds and spoken words, whereby the auditory input activates semantic object representations exerting top-down effects of expectation on the visual processing of objects.

## Acknowledgments

The project was funded by the European Union’s Horizon 2020 research and innovation programme under the Marie Sklodowska-Curie grant agreement No 659778, and by the European Research Council (ERC) under the European Union’s Horizon 2020 research and innovation programme (grant agreement No 725970). This manuscript reflects only the authors’ view and the Agency is not responsible for any use that may be made of the information it contains.

